# Cofilin-actin rod formation in neuronal processes after brain ischemia

**DOI:** 10.1101/331082

**Authors:** Seok Joon Won, Angela M. Minnella, Long Wu, Claire H. Eun, Eric Rome, Alisa E. Shaw, Paco S. Herson, James R. Bamburg, Raymond A. Swanson

## Abstract

Functional impairment after brain ischemia results in part from loss of neuronal spines and dendrites, independent of neuronal death. Cofilin-actin rods are covalently linked aggregates of colfilin-1 and actin that form in neuronal processes (neurites) under conditions of ATP depletion and oxidative stress, and which cause neurite degeneration if not disassembled. ATP depletion and oxidative stress occur with differing severity, duration, and time course in different ischemic conditions. Here we evaluated four mouse models of brain ischemia to define the conditions that drive formation of cofilin-actin rods. Three of the models provide early reperfusion: transient middle cerebral artery occlusion (MCAo), transient bilateral common carotid occlusion (CCAo), and cardiac arrest / cardiopulmonary resuscitation (CA/CPR). Early reperfusion restores ATP generating capacity, but also induces oxidative stress. The fourth model, photothrombotic cortical infarction, does not provide reperfusion. Cofilin-actin rods were formed in each of these models, but with differing patterns. Where acute reperfusion occurred, rod formation was maximal within 4 hours after reperfusion. Where infarction occurred, rods continued to form for at least 24 hours after ischemic onset, and extended into the adjacent non-ischemic tissue. Interventions that limit cofilin-actin rod formation may help to preserve integrity of neuronal processes in permanent ischemia.

## Introduction

Neuronal death contributes to functional disability after brain ischemia, and several mechanisms by which this occurs are now well established [1]. Loss of neuronal dendrites and spines may also contribute to functional disability [2-4], but far less is understood about these events. Ischemic injury can induce a loss of spines and dendrites even when the parent neuron survives [3, 4]. This subcellular level of injury is difficult to identify histologically or by standard noninvasive imaging modalities such as magnetic resonance imaging scans. To the extent that dendritic spines represent the anatomical correlate of experience, their loss would be expected to produce a transient, if not permanent functional deficit.

One mechanism by which ischemia could injure neurites is through sustained formation of cofilin-actin rods. Cofilin-1 is an actin-binding protein normally involved in the dynamic turnover of actin filaments (“actin treadmilling”) and other cell functions [5]. Cofilin-actin rods are formed when cofilin-1 is dephosphorylated by specific phosphatases, the dephosphorylated cofilin-1 binds to ADP-actin, and disulfide bonds are generated between the cofilin-1 molecules [6]. Oxidative stress, which in brain ischemia is induced through excitotoxic and reperfusion mechanisms, promotes cofilin-actin rod formation by forming intermolecular disulfide bonds between cofilin-1 cysteines 39 and 147 [6-8]. Oxidative stress may also promote rod formation by releasing a cofilin phosphatase, slingshot protein phosphatase-1L (SSH-1L) from its binding protein [9]. In addition, ATP depletion promotes cofilin-actin rod formation by increasing ADP-actin levels and by activating a second cofilin phosphatase, chronophin CIN [10].

Cofilin-actin rod formation halts local actin-dependent processes. This can have an immediate energy sparing effect because it interrupts local actin-dependent energy consuming processes such as synaptic vesicle docking and synaptic signaling, spine extension and contraction, dendritic transport, and mitochondrial trafficking [11, 12], and because actin turnover is itself a major source of neuronal energy expenditure [11, 13]. On the other hand, prolonged cessation of actin-mediated processes, such as organelle and protein trafficking, is not compatible with function or survival of neuronal processes [8, 14, 15].

Cofilin-actin rods can be imaged using antibodies to cofilin-1, as rod formation condenses the normally dispersed cofilin-1 into readily observable puncta and short rod-like segments. Neurons are the only cell type in brain in which cofilin-actin rods have been definitively localized. The rods form most readily in distal processes, especially dendrites, but can also form in cell bodies [8]. Cofilin-actin rods are biochemically distinct from the actin stress fibers that form in endothelial cells and muscle [16].

To our knowledge, there have been no prior published studies of cofilin-actin rod formation in brain ischemia. Ischemic brain injury (ischemic stroke) is broadly classified as either “transient ischemia” (or “ischemia-reperfusion”) when reflow occurs soon enough to preserve viability in at least a fraction of the ischemic tissue, or “permanent ischemia” when it does not. ATP depletion and oxidative stress occur with both types of ischemia, but with differing severity, duration, and time courses. Given that both ATP depletion and oxidative stress drive cofilin-actin rod formation, it is difficult to predict the patterns cofilin-actin rod formation that may occur in transient versus permanent ischemia. Accordingly, the aim of this study was to describe the time course and localization of cofilin-actin rod formation in four different mouse brain ischemia models. Three of these models involve reperfusion after variable intervals and somewhat differing anatomical regions of ischemia; transient middle cerebral artery occlusion (MCAo), transient bilateral common carotid occlusion (CCAo), and cardiac arrest / cardiopulmonary resuscitation (CA/CPF). The fourth model, photothrombotic cortical infarction, produces permanent ischemia without reperfusion. Our findings demonstrate robust and extensive formation of cofilin-actin rods in neuronal processes in each of these stroke models, and continued presence of cofilin-actin rods for at least 24 hours in the models that induce infarction.

## Materials and methods

### Animals

Studies were approved by the animal studies committees at the San Francisco Veterans Affairs Medical Center, and the University of Colorado, and were performed in accordance with the National Institutes of Health Guide for the Care and Use of Laboratory Animals. Results are reported in accordance with the ARRIVE guidelines [17]. Reagents were obtained from Sigma Aldrich, St. Louis, where not otherwise noted.

### Cell Culture studies

Mixed neuron-astrocyte cultures were prepared from the cortices of embryonic day 16 mice of both sexes and plated on poly-D-lysine coated glass coverslips. After 4 days in culture, 1.2μM cytosine arabinoside was added to culture media to suppress astrocyte proliferation. The cells were subsequently maintained in NeuroBasal medium (Gibco) containing 5% fetal bovine serum and 5 mM glucose, and used at day 13-15 in vitro. Experiments were initiated by exchanging the culture medium with a balanced salt solution (BSS): 1.2 mM CaCl_2_, 0.4 mM MgSO_4_, 5.3 mM KCl, 0.4 mM KH_2_PO_4_, 137 mM NaCl_2_, 0.3 mM NaHPO_4_, 5 mM glucose, and 10 mM 1,4-piperazinediethanesulfonate (PIPES) buffer, pH 7.3. Chemical oxygen-glucose deprivation (cOGD) was induced by adding 6 mM 2-deoxyglucose and 10 mM sodium azide to the BSS. After 20 minutes incubation at 37 °C, both the control BSS and cOGD BSS were replaced by fresh BSS, and after an additional 10 minutes incubation the cultures were fixed with phosphate-buffered 4% formaldehyde. Each experiment was performed on an independent culture preparation and included 4 replicates per condition.

### Transient focal cerebral ischemia

Adult (age 3-4 months) male C57BL6 mice were anesthetized with 2% isoflurane in 70% N_2_O/ balance O_2_ delivered through a ventilated nose cone. Rectal temperature was maintained at 37 ± 0.5 °C using a homeothermic blanket throughout the surgical procedure. Transient ischemia in the middle cerebral artery (MCA) distribution was induced by the Weinstein intraluminal filament method [18, 19]. The right common carotid artery was exposed through a midline incision, separated from the vagus nerve, and temporarily ligated. The distal part of external carotid artery was ligated, and a silicon-coated 6-0 monofilament (Doccol Corporation, Sharon, MA) was inserted through the proximal external common carotid artery stump and advanced along the internal carotid artery until the tip occluded the proximal stem of the MCA. Regional cerebral blood flow was monitored using Laser Doppler Flowmetry (Moor instruments Ltd, UK). After 30 minutes of occlusion, blood flow was restored by removal of the monofilament. The wound was sutured, bupivacaine (6 mg/kg) and meloxicam (7.5 mg/kg) were administered subcutaneously, and the mouse was moved to warmed recovery chamber until awake and ambulatory. For sham surgery, mice were subjected to the same procedure but without intraluminal suture insertion. Mice were excluded from the study if the measured cerebral blood flow was not reduced by more than 70% during vessel occlusion.

### Transient forebrain ischemia

Adult (age 3-4 months) male C57BL6 mice were anesthetized with 2% isoflurane in 70% N_2_O/ balance O_2_, delivered through a ventilated nose cone. Rectal temperature was maintained at 37 ± 0.5 °C by using a homeothermic blanket throughout the surgical procedure. Forebrain global ischemia was induced by occluding both common carotid arteries as described previously [20, 21]. The common carotid arteries (CCAs) were exposed bilaterally and isolated from the adjacent vagus nerves. Microvascular clips were applied to both CCAs. After 16 minutes of occlusion, clips were removed and patency of each CCA was confirmed by inspection under the surgical microscope. The wound was sutured, bupivacaine (6 mg/kg) and meloxicam (7.5 mg/kg) were administered subcutaneously, and the mouse was moved to warmed recovery chamber until awake and ambulatory. Sham-operated animals received the same neck skin incision but artery occlusion was not performed. Mice were excluded from analysis if reperfusion was not observed in both CCAs, or if mice developed seizures.

### Photothrombotic permanent ischemia

Adult (age 3-4 months) male C57BL6 mice were anesthetized with 2% isoflurane in 70% N_2_O/ balance O_2_, delivered through a ventilated nose cone. Rectal temperature was maintained at 37 ± 0.5 °C by using a homeothermic blanket throughout the surgical procedure. Photothrombotic permanent ischemia was induced by the Rose Bengal technique [22, 23]. A PE10 polyethylene catheter (Becton Dickinson, Franklin, NJ) was introduced into the femoral vein, and the head was immobilized in a stereotaxic frame. The skull was exposed by skin incision, and a 2 mm diameter fiber optic cable was place on the skull 1.0 mm anterior, and 1.5 mm lateral to bregma. Rose Bengal (Sigma-Aldrich, St Louis, MO; 20 mg/kg) dissolved in saline was infused through the venous cannula for 1 minute, and vascular occlusion was induced by 10 minutes illumination with white light (KL 1500 LCD, SCHOTT North America Inc., Southbridge, MA) through the fiber optic cable. Sham ischemia animals were injected with saline only but were otherwise treated identically. The incisions were sutured, bupivacaine (6 mg/kg) and meloxicam (7.5 mg/kg) were administered subcutaneously, and the mouse was moved to warmed recovery chamber until awake and ambulatory.

### Cardiac arrest / cardiopulmonary resuscitation

These studies were approved by the University of Colorado Institutional Animal Care and Use Committee. Adult (age 3-4 months) male C57BL6 mice were subjected to CA/CPR as previously described [24, 25]. Anesthesia was induced with 3% isoflurane and maintained with 1.5–2% isoflurane in oxygen enriched air via facemask. Temperature probes were inserted in the left ear and rectum to monitor head and body temperature simultaneously. A PE-10 catheter was inserted into the right internal jugular vein for drug administration. Needle electrodes were placed subcutaneously on the chest for continuous EKG monitoring. Animals were endotracheally intubated and connected to a mouse ventilator. Cardiac arrest was induced by injection of 50 μL KCl (0.5M) via the jugular catheter and confirmed by asystole on EKG. The endotracheal tube was disconnected and anesthesia stopped. During cardiac arrest, body temperature was allowed to spontaneously decrease to a minimum of 35.5 °C, and head temperature was maintained at 37.5 °C. Resuscitation was begun 8 min after induction of cardiac arrest by injection of 0.05–0.10 ml epinephrine solution (16 μg epinephrine/ml 0.9% saline), chest compressions, and ventilation with 100% oxygen at a respiratory rate of 200/min and 25% greater tidal volume. Chest compressions were stopped when spontaneous circulation was restored. Resuscitation was abandoned if spontaneous circulation was not restored within 2.5 min. Both catheters and temperature probes were then removed and skin wounds closed.

### Histology and immunocytochemistry

Anesthetized mice were perfused with cold saline (0.9% NaCl) followed by saline containing phosphate-buffered 4% formaldehyde (PFA). After post-fixation with 4% PFA for 24 hours, brains were immersed for another 24 hours in 20% sucrose for cryoprotection. The brains were then frozen and 40 μm coronal sections were prepared with a cryostat. The sections were permeablized with 95% methanol and 5% 0.1 M phosphate buffer for 15 minutes at -20 °C, then washed with phosphate-buffered saline. The fixed brain sections were pre-incubated in blocking buffer (2% donkey serum and 0.1% bovine serum albumin in 0.1 M phosphate buffer) at room temperature for 30 minutes, and then incubated with primary antibody overnight at 4 °C. After washing, the sections were incubated with fluorescent secondary antibody (Life Technologies Corporation, Grand Island, NY; 1:500) for one hour at room temperature. Where double labeling was additionally performed with cell-type specific antibodies, the sections were subsequently incubated with blocking buffer additionally containing 0.3% Triton X-100, and then incubated with those antibodies overnight at 4 °C followed by appropriate fluorescent secondary antibodies. Stained sections were mounted on glass slides in DAPI-containing mounting medium (Vector laboratories, Burlingame, CA). Control sections were prepared omitting either primary or secondary antibodies. Immunostaining of cell cultures was done identically except that incubation time with primary antibody was shortened to 6 hours.

Primary antibodies were obtained from the following sources and used at the designated dilutions: rabbit anti-cofilin-1, Cytoskeleton #ACFL02, Denver, CO (1:250); mouse anti-MAP2, Millipore #3418, Temecula, CA (1:1000); mouse anti-NeuN, Millipore #MAB377, Temecula, CA (1:1000); mouse anti-NF-H, BioLegend #SMI31, San Diego, CA (1:2000); goat anti-APP, Abcam #Ab2084, Cambridge, MA (1:400); mouse anti-CNPase, Promega #G346A, Madison, WI (1:800); mouse anti-GFAP, Millipore #MAB360, Temecula, CA (1:1000); goat anti-CD3, BioTechne #AF3628, Minneapolis, MN (1:200); rat anti-CD11b, BioRad # MCA711, Hercules, CA (1:250); mouse anti-mouse IgG, Vector Lab #AI-9200, Burlingame, CA (1:250). Mouse anti-cofilin1 #mAb22 (1:300) and rabbit anti- pospho-3-serine cofilin-1 # 4321 (1:2500) were provided by J. R. Bamburg, Colorado State University [26, 27].

### Image analysis of cofilin-actin rod formation

Confocal images were taken at three pre-determined locations on three sections from each animal. For photothrombotic stroke, this area was at the lateral edge of the ischemic cortical region. This region was difficult to define by visual inspection alone at the 1 and 4 hour time points, and for brains harvested at these time points the ischemic territory was identified by immunostaining for IgG, which is trapped in the vessels of the photo-thrombosed territory and not washed out during perfusion-fixation. The confocal images were z-stacked using the NIH ImageJ program and used for quantifying cofilin-1 immunoreactivity. Each processed image was prepared using a stack of 10, 1 μm Z-stage images (10 μm thickness). Background signal was defined as the signal level that excluded approximately 98% of pixels in a non-ischemic brain region from each section. This threshold was then applied to the ischemic region of interest, and the area of ischemia-induced cofilin signal was measured and expressed as percent of the image area (for brain sections) or percent of the MAP2 staining area (for cell cultures). Values for each image were averaged to provide an aggregate value for each brain. Both the person taking the photographs and the person analyzing the immunostaining were blinded to the treatment conditions.

### Image analysis of total cofilin-1 and ser-3-phospho-cofilin-1

Neuron cultures or brain sections were doubled labeled using rabbit antibody targeting serine-3-phospho-cofilin-1 [28] and mouse antibody targeting total cofilin-1 [29]. The signal from each antibody was assessed along 2 μm line segments oriented orthogonal to the axes of individual neurites. These segments were placed over sites of cofilin-actin rod formation in neurons treated with cOGD, and over non-rod-containing neurites in control cultures. (More than 95% of neurites had no rods in the control condition.) A background subtraction was made for each antibody signal using values obtained where no cells were present. The peak value of each of these determinations was used for aggregate statistical analysis. For assessments in brain sections, confocal images were taken in the peri-infarct regions of sections from 3 animals and a total of 29 cofilin-actin rods were selected for quantification using the same method as used for cultured cells, but without background subtraction.

### Statistics

All data are expressed as means ± s.e.m., with the “n” of each study defined as the number of mice or, for cell cultures, the number of independent experiments unless otherwise stated. Data were analyzed using one-way ANOVA and the Dunnett’s test for multiple comparisons against a common (control) group. Where only two groups were compared, the two-sided t-test was used.

## Results

We first validated cofilin-actin rod detection using a cell culture model of stroke, in which energy failure is induced by blocking glucose and oxygen utilization (chemical oxygen-glucose deprivation; cOGD). cOGD induced widespread formation of punctate and rod-shaped aggregates in neuronal processes that were recognized by rabbit antibody to cofilin-1 (Fig 1). These aggregates are characteristic of cofilin-actin rods [30, 31] were localized primarily to the neurites, though some were also observed in cell soma. Some neurites contained segments in which the rods were co-localized with microtubule-associated protein-2 (MAP2), whereas other segments showed an abrupt discontinuity of MAP2 at sites of rod formation (Fig 1). This pattern is in agreement with prior studies showing disruption of the neurite cytoskeleton by cofilin-actin rod formation [31-33]. Rare foci of cofilin-1 aggregation were also observed in control cultures that were treated with medium exchanges only.

**Fig 1.**
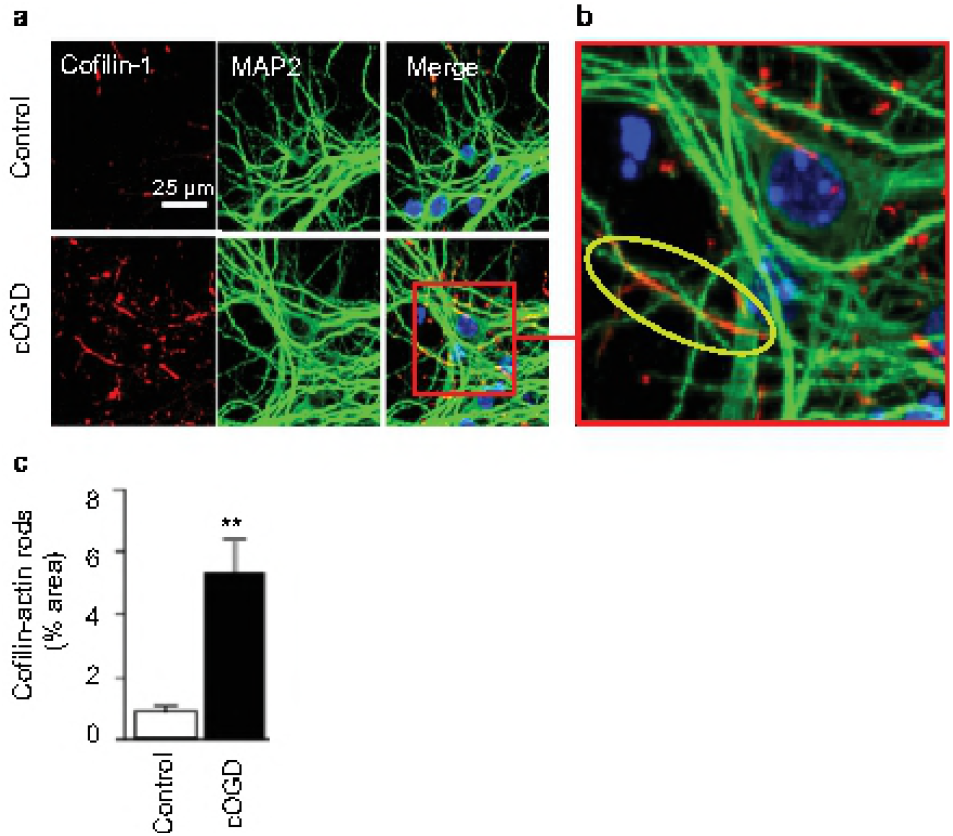
Oxygen-glucose deprivation induces formation of cofilin-actin rods in cultured neurons. **a**. Neurons are immunostained using antibodies to cofilin-1 (red) and MAP2 (green), with nuclei stained blue with DAPI. Cultures were treated for 30 minutes with chemical oxygen-glucose deprivation (cOGD) or medium exchanges only (control). **b**. Oval in enlarged view identifies a segment of neuronal process in which cofilin-actin rods have formed and MAP2 immunoreactivity is lost. C. Graph quantifies the relative area of cofilin staining in the control and cOGD conditions, expressed as percent of the area of MAP2 immunoreactivity. n = 4; **p < 0.01.

We next evaluated cofilin-actin rod formation in mice exposed to the transient MCAo model of stroke, using brains harvested 1, 4 and 24 hours after reperfusion. Transient MCAo produces a central core of near-zero blood flow, and a surrounding penumbral region with less severe ischemia [34]. Using this stroke model, we found no significant cofilin-actin rod formation at one-hour time post-reperfusion, but widespread formation at both the 4- and 24-hour time points (Fig 2). Sections from the 24-hour reperfusion group were then double-labeled for cofilin-1 and cell-type specific markers. The cofilin-1 immunostaining was restricted to cellular elements with the morphology and size of neuronal processes, and as in the cell culture studies some processes showed an abrupt discontinuity of cytoskeletal markers (MAP2 and NF-H) (Fig 3). The higher magnification views also show that many of the cofilin-actin rods that appear non-linear are composed of overlapping linear arrays. This is best seen in neurons co-labeled with the axonal marker amyloid precursor protein (APP), which is not a cytoskeletal element. By contrast, there was no co-localization between cofilin-1 and either the oligodendrocyte marker CNPase, the astrocyte cytoskeletal marker GFAP, the microglia/macrophage marker CD11b, or the endothelial cell marker CD31 (cluster of differentiation 31; also termed platelet endothelial cell adhesion molecule-1) (Fig 3).

**Fig 2.**
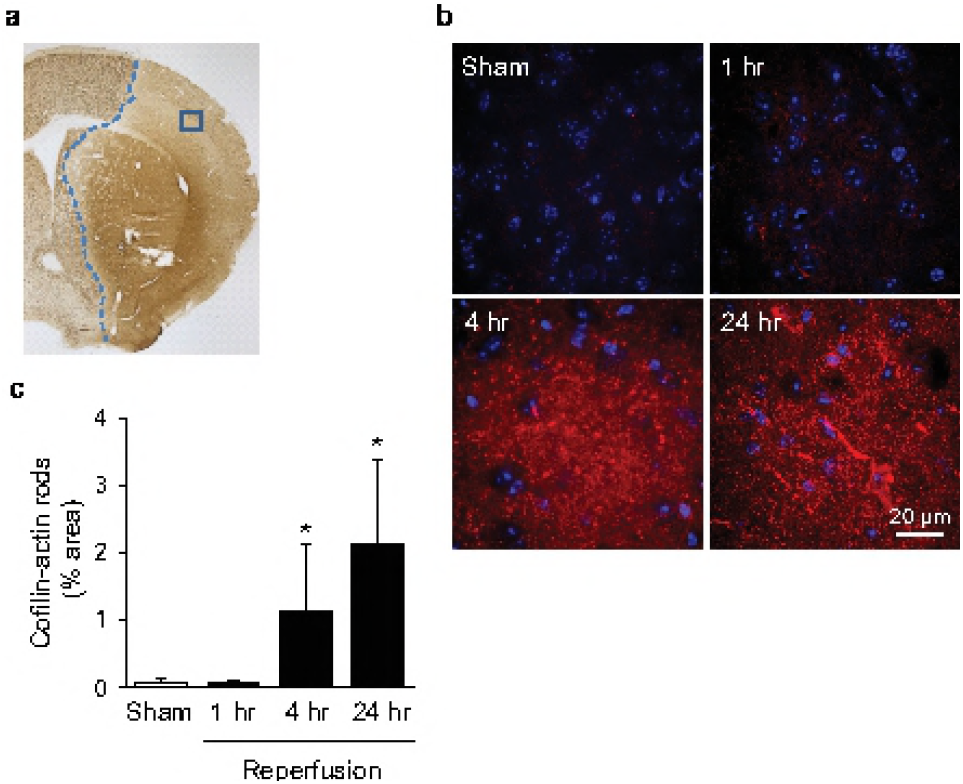
Cofilin-actin rod formation after transient focal ischemia. **a**. Lesioned area as identified by NeuN immunostaining in brain fixed 24 hours after a 30-minute MCA occlusion. Blue line marks the approximate medial edge of the lesion, and blue box shows the area photographed on each section after cofilin-1 immunostaining. **b**. Immunostaining for cofilin-1 (red) in the ischemic cortex at designated time points after reperfusion. Cell nuclei are stained blue. **c**. Graph quantifies area of cofilin staining at each time point, expressed as percent of the image area (n = 3-4; *p < 0.05 v. sham).

**Fig 3.**
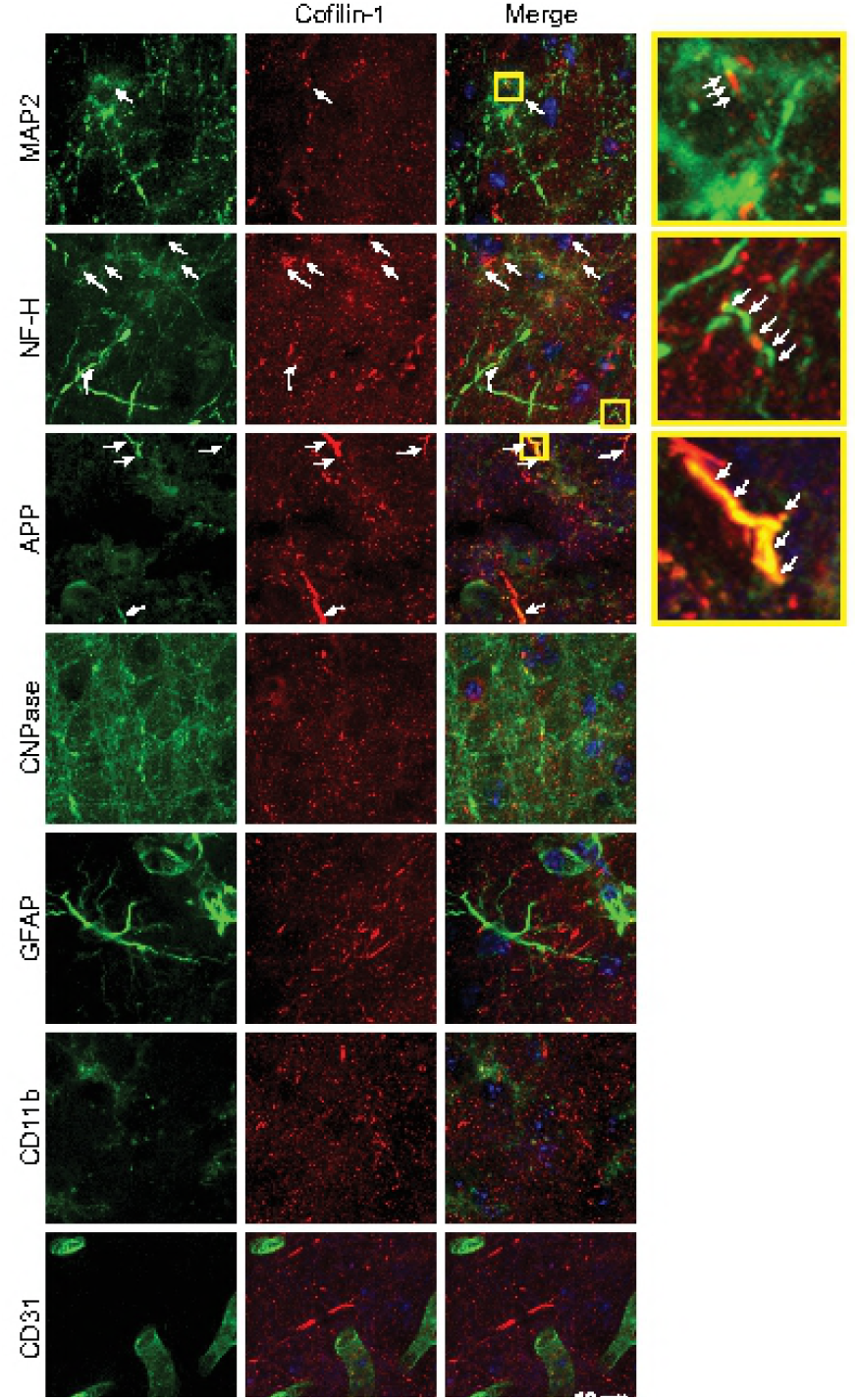
Cofilin-actin rods form in neurons. Panels show double immunostaining for cofilin-1 (red) and cell-type specific markers for neurons (MAP2 and NF-H), neuronal axons (APP), oligodendrocytes (CNPase), astrocytes (GFAP), microglia/macrophages (CD11b) and endothelial cells (CD31). Cell nuclei are stained blue. Yellow boxes denote the areas magnified in adjacent panels. Cofilin aggregates were observed in neuronal dendrites and axons (arrows) but not in other cell types. Brain sections were from ischemic cortex collected 24 hours after 30 minutes MCA occlusion and are representative of n = 4 mice.

We next evaluated cofilin-actin rod formation in the transient bilateral carotid artery occlusion (CCAo) of stroke. In mice, CCAo produces neuronal death preferentially in the forebrain cortex and in the hippocampal CA1 and dentate gyrus [20]. The neuronal death can be significantly attenuated by NMDA receptor antagonists or NADPH oxidase inhibition during reperfusion[21, 35, 36], indicating significant oxidative stress in these regions during ischemia-reperfusion. Immunostaining for cofilin-1 after CCAo showed rod shaped and punctate accumulation predominately in these same anatomical regions (Fig 4a). The morphology of cofilin-actin rods was similar to that observed after MCAo, but the time course was different. In CA1 and cortex, the extent of rod formation was several-fold greater than control by 1 hour after reperfusion, but less at 24 hours (Fig 4b). The pattern was different in the dentate gyrus, with peak signal observed at the zero time point (immediately before reperfusion, when ATP levels are likely lowest), and had resolved by 1 hour.

**Fig 4.**
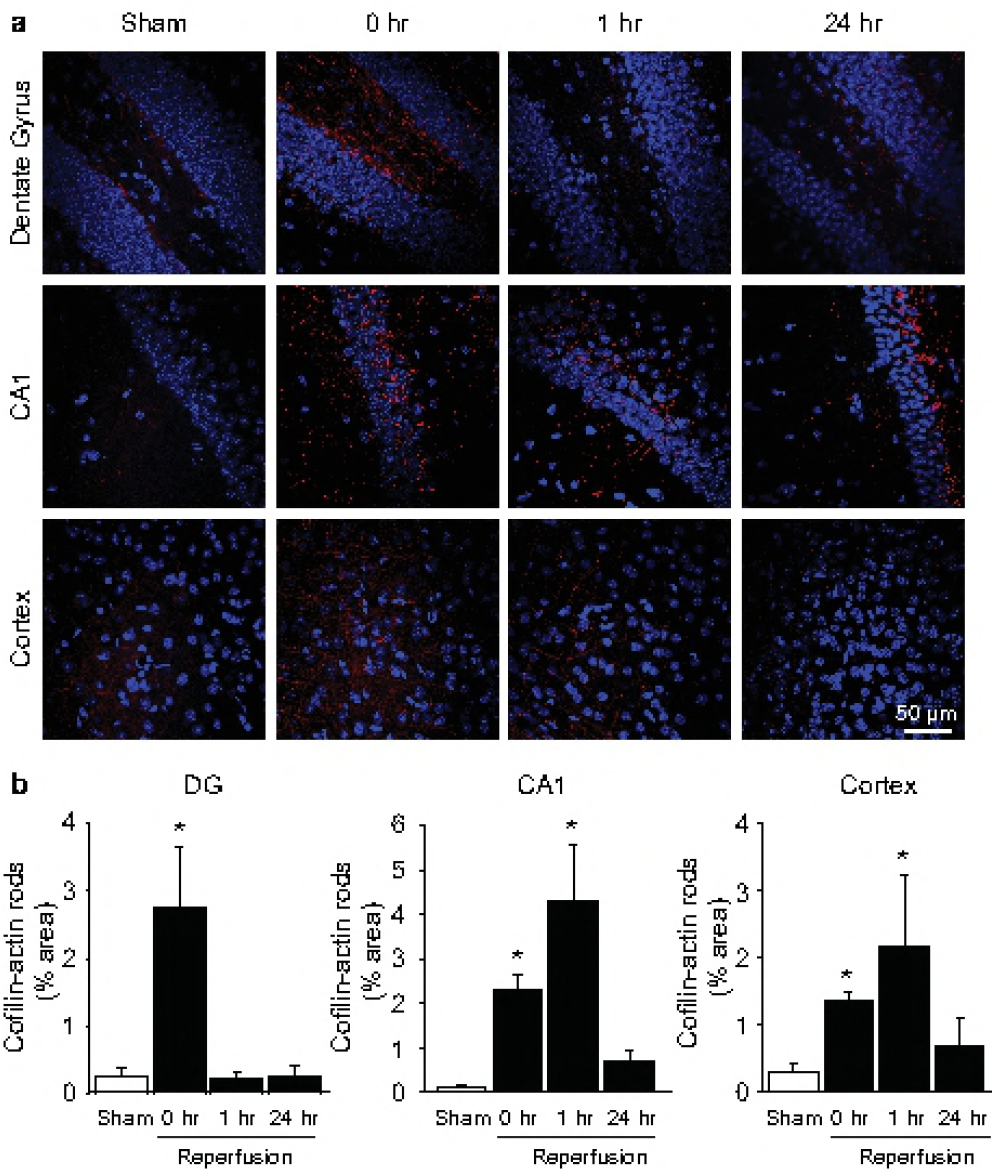
Cofilin-actin rod formation after transient common carotid artery occlusion (CCAo) **a**. Panels show immunostaining for cofilin-1 (red) in the hippocampal dentate gyrus (DG), hippocampal CA1, and cerebral cortex in sections obtained at designated time points after 16 minutes of bilateral common carotid artery occlusion. Cell nuclei are stained blue. **b**. Graphs show area of cofilin staining at each time point, expressed as percent of the image area. (n = 3; *p < 0.05 v. sham)

We additionally evaluated brains from mice subjected to cardiac arrest / cardiopulmonary resuscitation (CA/CPR) [24, 25]. CA/CPR renders the entire brain ischemic, but for a shorter interval than the CCAo model. Brains from CA/CPR mice showed negligible cofilin-actin rod formation in cortex (not shown), perhaps reflecting the shorter ischemic duration. Rod formation was observed in the in the hippocampal dentate gyrus and CA1 (Fig 5), though less extensively than in the CCAo mice.

**Fig 5.**
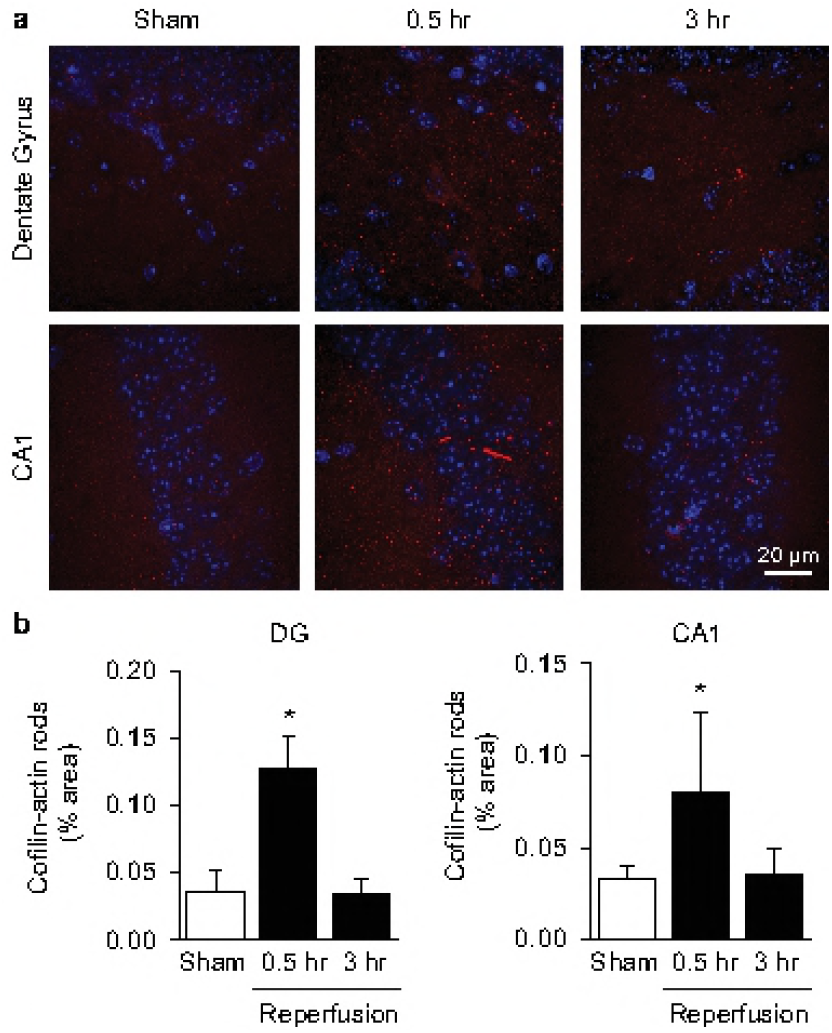
Cofilin-actin rod formation after cardiac arrest / cardiopulmonary resuscitation (CA/CPR) **a**. Panels show immunostaining for cofilin-1 (red) in the hippocampal dentate gyrus (DG) and hippocampal CA1 in sections obtained at 30 minutes or 3 hours after 8 minutes of cardiac arrest. Cell nuclei are stained blue. **b**. Graphs show area of cofilin staining at each time point, expressed as percent of the image area. (n = 3; *p < 0.05 v. sham)

Most clinical strokes are caused by emboli that persist for many hours or days, and hence may better be mimicked by permanent ischemia animal models that do not provide early reperfusion. To evaluate cofilin-actin rod formation in permanent ischemia, we used a photothrombotic method to induce infarction in a defined region of mouse cortex (Fig 6a). Cofilin-actin rods were identified primarily on the border region of the resulting infarct, and not in the infarct core (Fig 6b,c). In contrast to the stroke models with early reperfusion, rod formation was observed to increase between 1 and 24 hours after ischemia onset, and expand out into the non-ischemic tissue in which normal cell nuclei were apparent (Fig 6d).

**Fig 6.**
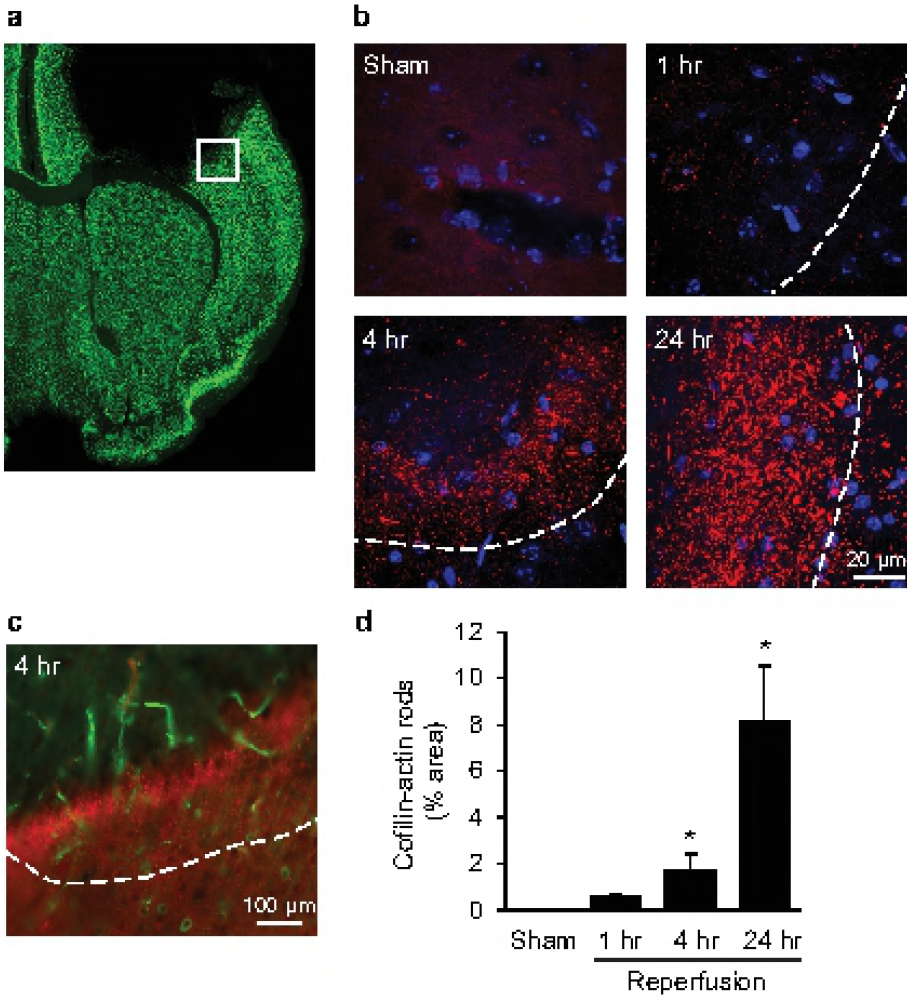
Cofilin-actin rod formation in photothrombotic permanent ischemia. **a**. Representative brain section immunostained for NeuN, showing location (white box) of photographs obtained. **b**. Cofilin-1 immunostaining (red) in the peri-infarct cortex at the designated time points after photothrombotic vascular occlusion. White line denotes infarct edge, and cell nuclei are stained blue. **c**. Lower power view of infarct edge, showing IgG retention in blood vessels (green) used to define the area of photothrombosis. Cofilin-actin rods (red) are formed at the edge of the infarct and extend into the non-ischemic tissue, but are not formed in the core of the photothrombosed area. **d**. Graph shows area of cofilin staining at each time point, expressed as percent of the image area. (n = 3-4; *p< 0.05 v. sham).

Cofilin-actin rod formation requires de-phosphorylation of the serine-3 residue of cofilin-1. Thus, to further confirm the nature of the aggregates detected by cofilin-1 immunostaining, we used neuron cultures exposed to cOGD and then probed these with antibodies that target either total cofilin-1 or specifically ser-3-phosphocofilin-1 [26, 28, 29]. The antibody recognizing 3-ser-phosphocofilin was generated in rabbit, and consequently a mouse antibody to total cofilin-1 was used to perform the double labeling. As shown Fig 7a, the mouse antibody to total cofilin-1 showed the same pattern of rod formation after cOGD as observed with the rabbit antibody to total cofilin-1 in Fig 1. Double labeling for ser-3-phosphocofilin-1 showed globally reduced immunoreactivity in cOGD-treated neurons, along with a striking reduction in the ratio of ser-3-phosphocofilin-1: total cofilin-1 at sites of cofilin-actin rod formation. This double-labeling approach was then used to similarly assess post-ischemic brain sections. As shown in Fig 7d,e a reduction in the ratio of ser-3-phosphocofilin-1: total cofilin-1 was also observed at sites of ischemia-induced cofilin-actin rod formation *in vivo*. The magnitude of this reduction appeared to be less than in the cell cultures, possibly because of signal from tissue above and below the selected neurites (despite confocal imaging) or partial re-phosphorylation of free cofilin-1 in the 24 hour interval between ischemia and brain harvest.

**Fig 7.**
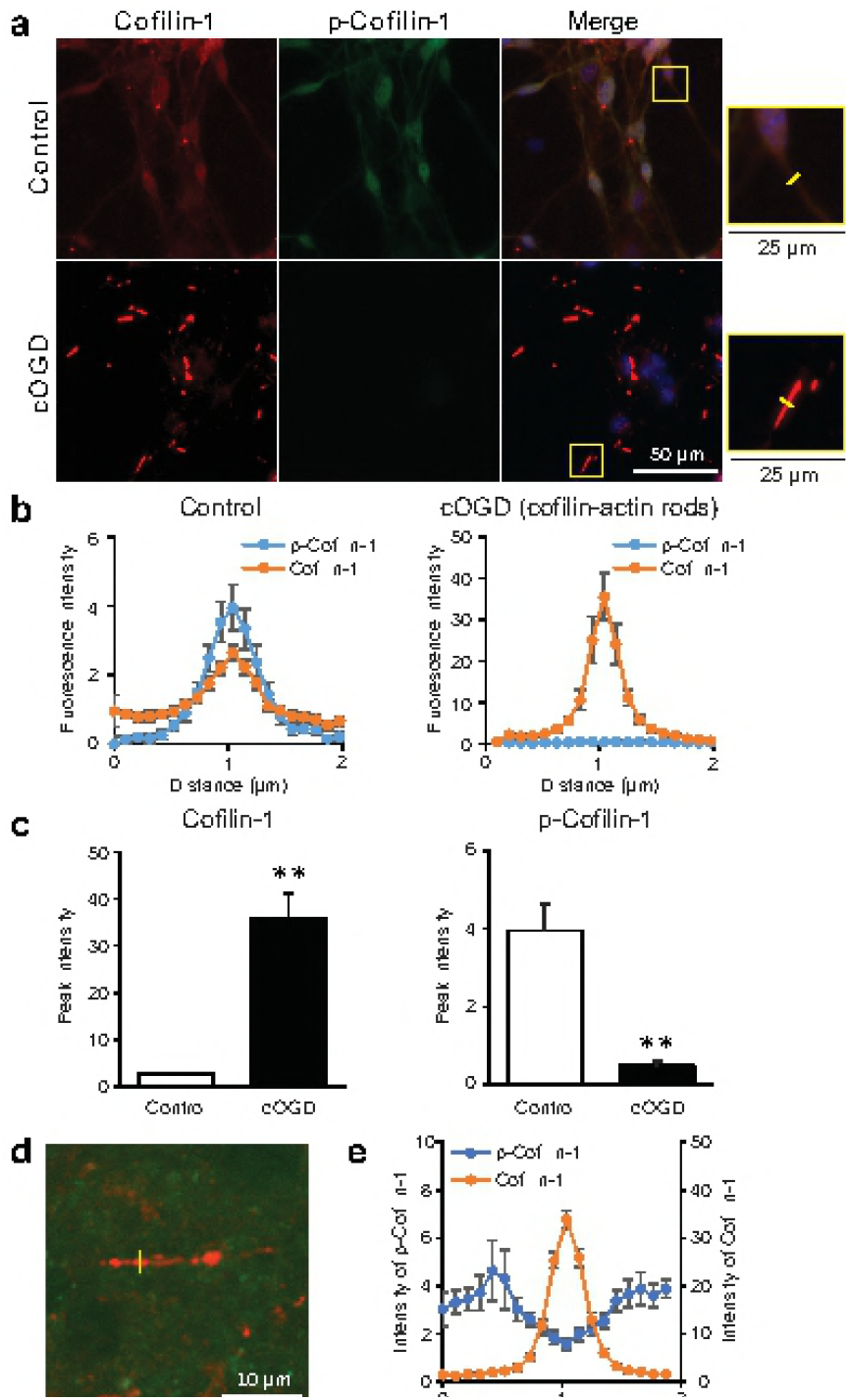
Reciprocal changes in cofilin-1 and phospho-cofilin-1 during rod formation. **a.**Immunostaining for total cofilin-1 (red) and serine-3-phospho-cofilin-1 (p-cofilin-1; green) in primary neuron cultures following 1 hour of chemical oxygen-glucose deprivation (cOGD) or medium exchange only (Control). Absolute intensity of the total cofilin-1 signal was in all cases much greater than the p-cofilin-1 signal, and in order to show double labeling the photograph settings were adjusted so that brightness of the cofilin-1 and p-cofilin-1 signals were comparable under control conditions. Nuclei are stained blue with DAPI in the merged image. Magnified views show neuronal processes. **b**. Fluorescence intensity of total cofilin-1 and p-cofilin-1 as measured along line segments drawn orthogonal to neuronal processes, as pictured in yellow in the magnified views of (**a**). Measurements were made in 10 neurites in cOGD – treated cultures that contained cofilin-actin rods, and 10 neurites in control cultures (that did not have rods). Note different Y-axis values in the two graphs. **c**. Aggregated peak fluorescence intensities in neurites measured as in (**b**). Note different Y-axis scales in the two graphs. n = 10 neurites; **p< 0.01. **d**. Confocal image from mouse brain fixed 24 hours after a 30-minute MCA occlusion,with double labeling for total cofilin-1 (red) and p-cofilin-1 (green). **e**. Measurement of total cofilin-1 and p-cofilin-1 immunoreactivity across cofilin-actin rods in post-ischemic brain sections shows reduced p-cofilin-1 signal at the sites of cofilin-actin rod formation. n = 29 neurites analyzed (from 3 brains).

## Discussion

Our primary aim was to describe and compare cofilin-actin rod formation in established brain ischemia models that produce differing components of reperfusion injury. For technical reasons it was not possible to evaluate uniform time points in each ischemia model, but nevertheless several distinct patterns emerge from these studies. First, the two models with brief ischemic intervals, CCAo and cardiac arrest, both induced rods preferentially in areas known to exhibit high levels of oxidative stress with ischemia-reperfusion; the hippocampal CA1 and dentate gyrus. In both of these models rod formation was maximal within 3 hours of reperfusion and had largely resolved at later time points. By contrast, the two models causing frank infarction (MCAo and photothrombosis) both produced a progressive increase in rod formation over the 24 hour observation interval. The photothrombotic model uniquely permits clear delineation of the ischemic border, and here rod formation was found to extend into the non-ischemic territory at the 24 hour time point. Importantly, rod formation was not observed in the core of the photothrombotic stroke, indicating that it is not simply an inevitable aspect of neuronal death. Notably, in all four stroke models cofilin-actin rod formation was significant long before neuronal degeneration occurs.

Cell culture studies have shown that oxidative stress promotes cofilin-actin rod formation through effects on cofilin phosphorylation state and intermolecular disulfide bonding [6-8]. Our findings indirectly support oxidative stress as a major factor promoting rod formation in brain ischemia. Oxidative stress is induced in transient ischemia by reperfusion of oxygenated blood, and at the margins of permanent ischemia by excitotoxicity. Here, brain ischemia induced rod formation in regions of reperfusion, and at the edge (but not core) of the permanent photothrombotic lesions.

Support for a specifically neuronal localization of the cofilin-actin rods in these studies is provided by the size and morphology of the cofilin-immunoreactive structures, the lack of co-localization with other cell-type specific markers, the presence of cofilin-actin rods in processes contiguous with neuron-specific markers, and their similar appearance to those produced in neuronal cultures treated with cOGD. The anti-cofilin antibodies used for these studies have been extensively characterized in neurons and other cell types [26-29]. The antibody against phosphoserine 3 of cofilin-1 also recognizes the phosphorylated serine 3 of the closely related protein ADF (actin depolymerizing protein); but ADF is present low levels relative to cofilin-1 in mouse neurons cells [26]. The culture studies presented here were performed primarily to validate the reagents used for cofilin-actin rod detection, but they additionally show MAP2-postive neurites containing discrete segments of cofilin-actin rod formation and disrupted MAP2 signal. This same pattern was observed *in vivo* following ischemia, further supporting a specifically neuronal localization of cofilin-actin rod formation. The reason why rod formation causes cytoskeletal disruption is not known, but may be related to actin sequestration in rods and cessation of normal transport along neuronal processes [14, 31, 32, 37].

Evidence suggests that rod formation reduces energy expenditure over short time intervals, but leads to neurite degeneration if sustained [6-8]. Given that energy failure is a primary initiator ischemic cell death, the formation of cofilin-actin rods could have either beneficial or deleterious effects on neurite survival depending upon ischemia duration and other factors. It is also possible that rod formation in neurites contributes to the survival (or death) of the parent neuron either through energy-related processes or other recognized mechanisms [38].

The exploratory design of this study required some limitations. In order to evaluate multiple time points in each of four stroke models, we used small numbers of mice at each time point, and used only adult male mice to minimize variability within each treatment condition. It is possible that age or sex may influence rod formation or resolution in ways that differ from the patterns observed here. Most importantly, these descriptive findings do not establish whether cofilin-actin rod formation has a causal effect on cell or neurite survival, or on long term functional outcomes. Tests of these potential cause-effect relationships will require studies in which rod formation is suppressed or promoted. Reagents are now becoming available to this end [39-41], and the observations presented here should help provide a framework for such efforts.

## Acknowledgements

This work was supported by the National Institutes of Health (Grant number R01AG049668, JRB; Grant number R01NS081149, RAS), and by the Dept. of Veterans Affairs. These sponsors were not involved in study design, data analysis, decision to publish, or other aspects of this work. The author(s) declared no potential conflicts of interest with respect to the research, authorship, and/or publication of this article. S.J.W., A.M.M. and R.A.S. designed research; S.J.W., A.M.M., L.W., C.H.E., A.E.S. and E.R. performed the experiments; P.S.H. provided mouse brains after cardiac arrest / cardiopulmonary resuscitation; J.R.B. provided a critical reagents and methods; S.J.W., A.M.M., L.W., C.H.E. and E.R. analyzed data; S.J.W., R.A.S, and J.R.B. wrote the manuscript.

